# Robust 3D Bloch-Siegert based B_1_^+^ mapping using Multi-Echo General Linear Modelling

**DOI:** 10.1101/550509

**Authors:** Nadège Corbin, Julio Acosta-Cabronero, Shaihan J. Malik, Martina F. Callaghan

## Abstract

**Purpose:** Quantitative MRI applications, such as mapping the T_1_ time of tissue, puts high demands on the accuracy and precision of transmit field (B_1_^+^) estimation. A candidate approach to satisfy these requirements exploits the difference in phase induced by the Bloch-Siegert Shift (BSS) of two acquisitions with opposite off-resonance frequency RF pulses. Interleaving these RF pulses ensures robustness to motion and scanner drifts, however, here we demonstrate that doing so also introduces a bias in the B_1_^+^ estimates.

**Methods:** It is shown here via simulation and experiments that the amplitude of the error depends on MR pulse sequence parameters, such as TR and RF spoiling increment, but more problematically, on the intrinsic properties, T_1_ and T_2_, of the investigated tissue. To solve these problems, a new approach to BSS-based B_1_^+^ estimation that uses a multi-echo acquisition and a general linear model (GLM) to estimate the correct BSS-induced phase is presented.

**Results:** In line with simulations, phantom and in-vivo experiments confirmed that the GLM-based method removed the dependency on tissue properties and pulse sequence settings.

**Conclusion:** The GLM-based method is recommended as a more accurate approach to BSS-based B_1_^+^ mapping.

## Introduction

Knowledge of the spatial distribution of the radiofrequency transmit field (B_1_^+^) is crucial to many MRI applications. Moderate accuracy may suffice when setting transmitter gain^1^ or calibrating multi-channel systems^2^. However, very high accuracy and precision are required for many quantitative MRI applications, e.g. mapping the longitudinal relaxation rate to characterise cortical myelination^3,4^.

Phase-based methods may be preferred as they are theoretically insensitive to T_1_ relaxation effects which often bias magnitude-based methods, especially at short TR. In the Bloch-Siegert (BS)^5,6^ approach, an off-resonance RF pulse leads to the Bloch-Siegert frequency shift (BSS), and an associated phase accumulation, which is proportional to the square of the pulse amplitude thereby encoding the B_1_^+^ field. This technique performed favourably in a recent review of the accuracy, precision and practicality of a range of prominent B_1_^+^ mapping techniques^7^, and has been shown to be less sensitive to *B*_0_ inhomogeneities and chemical shifts^8^ than other phase-based methods.

The BSS technique is flexible and can be integrated into a multitude of pulse sequences, such as 2D^9^ or 3D gradient echo (GRE)^8,10,11^, Interleaved EPI, spiral GRE^12^ and spin echo^13–16^ acquisitions. The BSS technique, as typically implemented, requires two acquisitions with opposite off-resonance frequencies. By subtracting the two phase images, the BSS effect is enhanced and unrelated phase components are removed, e.g. phase accumulated across echo time (TE) due to B0 inhomogeneity or chemical shifts, due to eddy currents, or any initial phase due to the transmitting and receiving coils. This subtraction also has the advantage of removing the effect of B_0_ inhomogeneity on the BSS, up to first order^9^. These two acquisitions can either be played out sequentially or by interleaving the opposite, off-resonance frequencies. Previous reports have shown the interleaved approach to be more robust to motion^17^ and magnetic field drift^10^.

In this work, we focus on a 3D spoiled GRE implementation and demonstrate, through both simulations and experiments, that alternating the sign of the off-resonance frequency from shot to shot in an interleaved fashion disturbs the steady-state and introduces an additional phase difference between the two acquisitions, especially at short TR. We show that this additional phase difference leads to bias in the B_1_^+^ map that depends on the relaxation parameters of the studied tissue, the specifics of the RF spoiling regime and the actual B_1_^+^ amplitude. We additionally propose and validate a modified BSS-based approach that removes these dependencies. The solution consists of a multi-echo acquisition in which several echoes are acquired before and after the BS pulse and modelling the phase evolution with a General Linear Model (GLM). We demonstrate that this GLM-based approach to isolating the BSS phase allows the interleaved approach to be used without introducing any error, extending the acquisition time, increasing the SAR or reducing the sensitivity.

## Theory

In line with Sacolick *et al*.^9^, and as detailed in the Supporting Information, the BS phase introduced by an RF pulse with peak amplitude 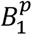 and normalised shape 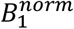, is proportional to the square of the peak pulse amplitude:

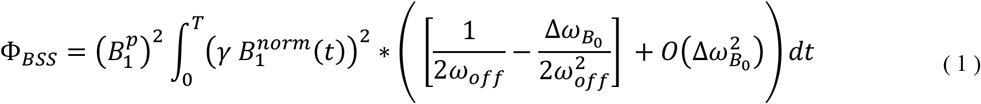

*ω_off_* is the off-resonance frequency of the pulse and Δ*ω*_*B*_0__ is the local field inhomogeneity, both in Hz.

### The classic method: isolating the BSS phase by subtraction

The classic approach to BSS-based B_1_^+^ mapping consists of acquiring two datasets with BS pulses of opposite off-resonance frequency (i.e. +*ω_off_* and – *ω_off_*). The phase difference between these is:

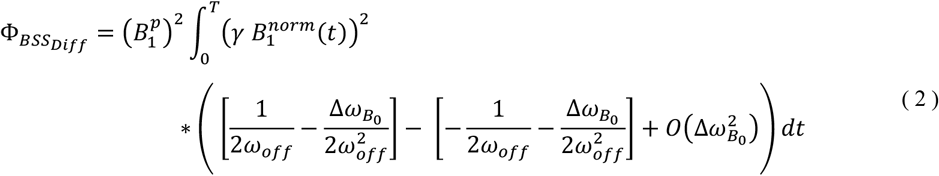

Since the first order terms that depend on Δ*ω*_*B*_0__ cancel, this expression simplifies to:

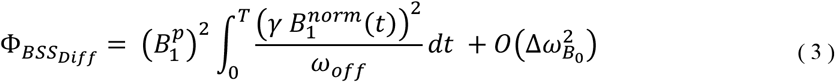

Previously this subtraction was assumed to also remove any phase accumulated from other sources, such as eddy-currents, transmit/receive-related phase offsets, chemical shifts and local B_0_ inhomogeneities. However, crucially, this is only true if the additional phase components are identical for each of the off-resonance frequencies. If this assumption is violated, the B_1_^+^ estimate will be erroneous.

### The GLM method: isolating the BSS phase by modelling a multi-echo acquisition

We propose an alternative to the classic BSS approach that computes accurate B_1_^+^ maps even if conditions vary between the two off-resonance frequency acquisitions. This approach relies on two novel features:

#### 1. A dual-offset multi-echo sequence

In the modified BSS-based B_1_^+^ mapping sequence (Fig.1), multiple echoes are acquired after one excitation pulse. Two echoes, after the BS pulse, have previously been used^18^ to concurrently compute the B_0_ inhomogeneity, whereas here multiple echoes are acquired before and after the BS pulse. As in the classic method, a second acquisition is performed with the opposite off-resonance frequency, either sequentially or interleaved.

**Figure 1:**
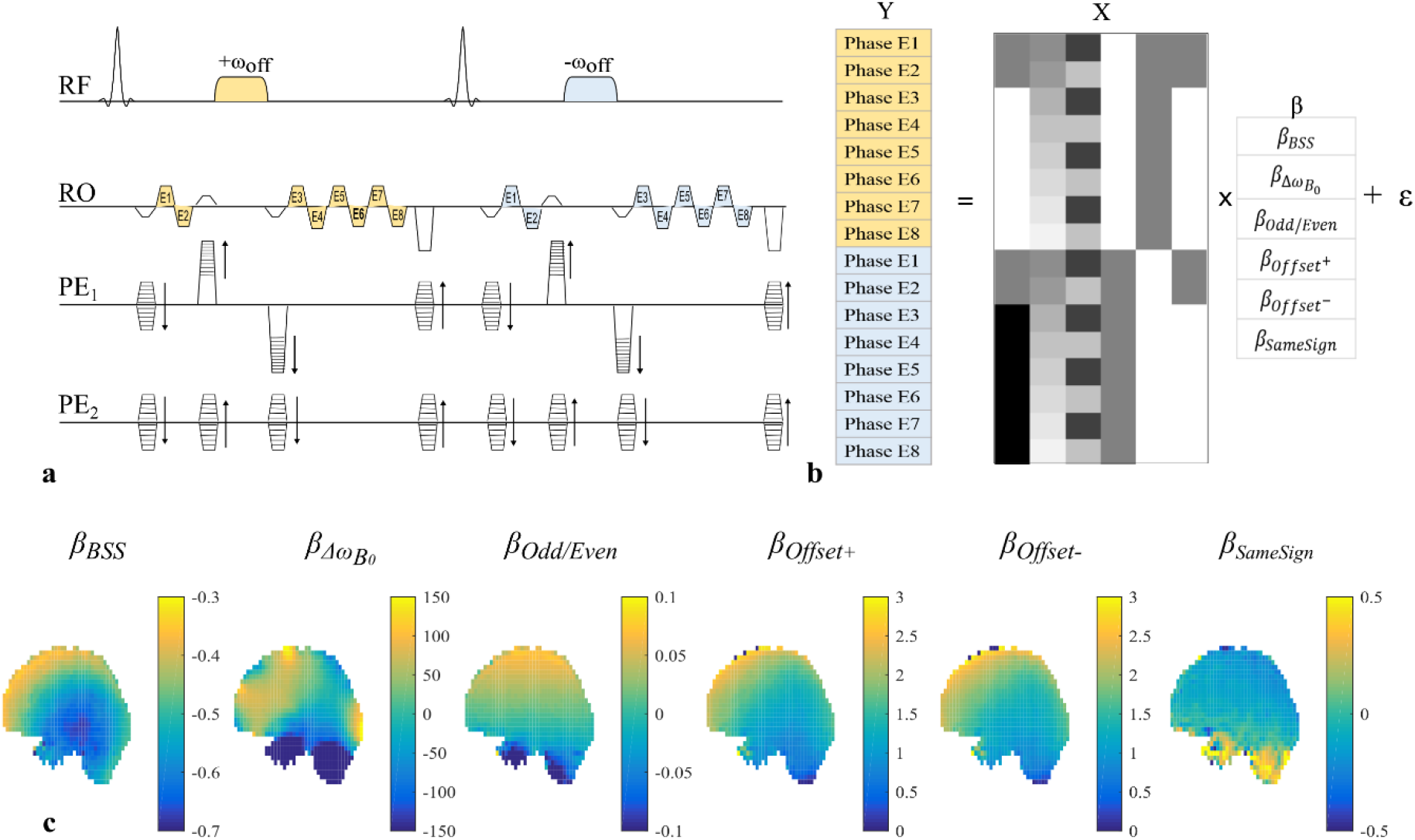
The sequence diagram of the modified 3D multi-echo gradient echo for BSS-based mapping is illustrated in (a). Two echoes are acquired prior to, and six echoes after, the Bloch-Siegert pulse, which is flanked by crushers in one phase-encoding direction (PE_1_) to destroy any inadvertent on-resonance excitation and minimize dependence on excitation flip angle. The gradients on each axis are balanced prior to the BS pulse. In this example, 8 echoes are acquired for each off-resonance frequency with two before and 6 after the BS pulse, resulting in a total of 16 phase images from which the B1+ efficiency is mapped. A GLM is used to model the phase variation across echo times (b). Typical maps of model coefficients obtained in vivo (c) exemplify the phase accrued due to the Bloch-Siegert shift (β_BSS_), B0 field inhomogeneity, (β_Δω_B_0___), alternating readout (RO) polarity, (β_Odd/Even_), initial phase offsets specific to the off-resonance frequency of the pulse, (β_Offset_+, β_Offset_-), and any additional phase due to the presence of the block of crushers and the BS pulse that is independent of the sign of the off-resonance frequency of the pulse, (β_SameSign_).

#### 2. A general linear model

The GLM approach models the phase data, Y, as multiple linear sources of phase accumulation over time: *Y* = *Xβ* + *ϵ*.

Each row of the model matrix X, corresponds to a single echo. The following explanatory variables model distinct sources of phase evolution in separate columns:

- *X_BSS_*: Models the phase accumulated due to the BS pulse, specifically the first term of the sum in Eq.1 (only). This phase should only be present after the BS pulse, and should change sign with the off-resonance frequency. The regressor consists of zeros for echoes before the BS pulse and either 1 or −1 afterwards depending on the BS pulse frequency.
- *X_Δω_B_0___*: Models the phase accumulated due to local *B*_0_ inhomogeneities. Regressor values are echo times.
- *X_Odd/Even_*: Models the phase difference between odd and even echoes due to eddy currents generated by the bipolar readout gradients. *X_Odd/Even_* is 1 for odd echoes and −1 for even echoes.
- *X*_*Offset*^+^_ and *X*_*Offset*^−^_: Models the initial phase offset of the acquisitions with positive and negative off-resonance frequencies respectively. *X*_*Offset*^+^_ is 1 for all echoes from a TR with a BS pulse with positive off-resonance frequency, and 0 for all echoes from a TR with a BS pulse with negative off-resonance frequency; vice versa for *X*_*Offset*^−^_.
- *X_SameSign_*: Models phase consistently accumulated during the BS pulse and the crushers, regardless of the sign of the BS pulse’s off-resonance frequency. This includes the second term in Eq. 1 and, for example, any phase due to eddy currents generated by the crushers. *X_SameSign_* is 0 for echoes prior to the BS pulse and 1 for echoes after the BS pulse, regardless of the pulses off-resonance frequency.

The parameters *β* constitute the regression coefficients and are estimated voxel-wise via the Weighted Least Square approach: 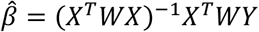. W is a diagonal weighting matrix in which the weights are the magnitude of the echoes. *β*_BSS_, the parameter of interest, is considered an estimate of Φ*_BSS_*. *ϵ* is an error term.

## Methods

### Numerical simulations

Simulations were used to evaluate the difference, both before and after the BS pulse, between two acquisitions with opposite off-resonance frequencies. Any difference present at this point would introduce an error in B_1_^+^ estimated with the classic method. A typical GRE acquisition was simulated by a series of matrix operations (described in Supporting Information). Several configurations were investigated:

❖ *Effect of RF Spoiling increment* RF spoiling modifies the phase of the excitation pulse and the BS pulse (*ϕ*) so that the phase difference between successive TR periods increases linearly by a constant amount *ϕ_BaseInc_*. The impact of the RF spoiling increment was investigated by changing *ϕ_BaseInc_* from 0° to 180° in 10° steps. A phase increment of 0 corresponds to no RF spoiling (i.e. *ϕ* = 0 for all pulses).
❖ *Sequential or Interleaved* For an interleaved acquisition, the sign of *ω_off_* was alternated between successive BS pulses. For sequential acquisitions, the sign was switched after half the total number of pulses 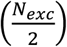, at which point RF spoiling was also reset.
❖ *Effect of TR, T_1_, T_2_ and 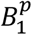* Extreme values were used to test the impact of sequence parameters, 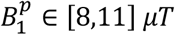, *TR* ∈ [35,100] *ms*, relaxation parameters, *T_1_* ∈ [550,1350] *ms* and *T_2_* ∈ [70,100] *ms* on the B_1_^+^ estimate.

The estimated B_1_^+^ amplitude was calculated as follows having isolated the first term (only) of Eq.1:

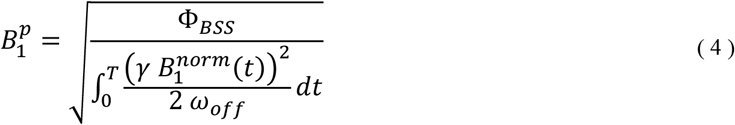

For the Classic method, Φ*_BSS_* was taken to be half the difference in phase accumulated after the BS pulse of the acquisitions with opposite off-resonance frequency, and termed 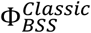 (see Supporting Information).

For the GLM method, Φ*_BSS_* was taken to be the mean absolute phase accrued during the BS pulses of these, and termed 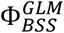 (see Supporting Information).

### MRI measurements

#### ∎ MR pulse sequence

Measurements were performed at 3T (Siemens Prisma) using a body coil for transmission and a 32-channel head coil for signal reception using an in-house MR pulse sequence (Fig.1). A Fermi pulse of duration T=2ms imparted the Bloch-Siegert shift after the second echo. Acquiring two echoes before the BS pulse served to minimize the correlation of the *X_BSS_* regressor with other regressors while maintaining a reasonable TE for the echo after the BS pulse. The encoding gradients on all axes were balanced immediately prior to the BS pulse, to ensure the same dephasing state for the magnetisation across TRs, and played again just after. Crusher gradients were played out either side of the BS pulse, concurrently with the balancing/phase-encoding gradients (on PE_1_), to crush any undesired on-resonance excitation and to minimise any dependence on the excitation flip angle. As demonstrated by Duan *et al*.^18^, perfect dephasing of the transverse magnetization before the BS pulse is required to fully remove any such dependence. Sensitivity to non-ideal conditions is reduced by using a high crusher moment since the greater the dephasing, the smaller the dependence on the excitation flip angle. Therefore a relatively large crusher moment^19^, designed to generate a theoretical dephasing of 6*π* rad across a voxel, was used. An excitation flip angle of 15°, corresponding to the Ernst angle for a TR of 35 ms and a T_1_ of 1000ms, was chosen to maximise the precision of the B_1_^+^ mapping^18^. Although this may be somewhat sub-optimal from a precision perspective for the phantom acquisitions and the long TR in vivo acquisitions, it was used for all acquisitions to ensure consistency.

For sequential acquisitions the positive and negative off-resonance frequency pulses were played out in consecutive blocks. In the interleaved case the off-resonance frequency was alternated across successive TR periods. To achieve steady state, 200 dummy cycles were executed before each block in the sequential case, 400 cycles were used at the outset of the interleaved case (to match acquisition times), and in both cases a spoiler gradient, set to reach a dephasing of 6*π* at the end of the TR, was applied in the readout direction after the last echo. The RF spoiling was reset at the end of the first block in the sequential case.

#### ∎ B_1_^+^ map estimation

All data, including B_1_^+^ maps, were reconstructed in real time using in-house code implemented in Gadgetron^20^. An apodizing filter was applied to the raw k-space data along each dimension to minimize ringing artefacts. After Fourier transformation the images were adaptively combined across coil elements^21^.

Two B_1_^+^ maps were computed for each dataset using the Classic and GLM methods respectively. For the Classic method, the phase of the third echoes, which were acquired after the positive and negative off-resonance frequency BS pulses, were subtracted to estimate the BSS phase: Φ*_BSS_* = Φ*_Diff_*/2. For the GLM method, the phase images for each off-resonance frequency were temporally unwrapped, by spatially unwrapping the differences between successive echo pairs^22^, and cumulatively adding these to the phase of the first echo. The phase images were subsequently used to estimate the parameters of the GLM (Fig.1). The BSS phase Φ*_BSS_* was captured by the first regressor *X_BSS_* of the model matrix such that Φ*_BSS_* = β_BSS_.

For both methods, 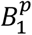 was computed on a voxel-wise basis using Eq.4.

#### ∎ Phantom experiments

B_1_^+^ maps were acquired on an FBIRN gel phantom^23^. Experiments 1 and 2 used both the Classic and GLM processing methods to construct B_1_^+^ maps. B_1_^+^ errors were quantified as the percent difference of the estimated B_1_^+^ with respect to a reference B_1_^+^ map.

##### Phantom Experiment 1: Comparing processing approaches

This experiment probed the impact of the processing method as a function of the RF spoiling increment, *ϕ_BaseInc_* ∈ [0; 10; 20; 50; 60; 70; 90; 110; 120; 130; 160; 170; 180]). The interleaved acquisition scheme was used for all scans. B_1_^+^ maps were reconstructed with the Classic and GLM methods. The voxel-wise difference between the two B_1_^+^ maps (relative to the GLM method) was computed, and summarised by the mean and standard deviation across the phantom. A TR of 35 ms was used.

##### Phantom Experiment 2: Comparing sequential and interleaved approaches

A reference B_1_^+^ map was obtained, via the Classic method, using a sequential acquisition with ϕ*_BaseInc_* = 90°. Four interleaved acquisitions were performed with:

1. No RF spoiling, TR=35ms
2. ϕ*_BaseInc_* = 120°, TR=35ms
3. ϕ*_BaseInc_* = 90°, TR=35ms
4. ϕ*_BaseInc_* = 90°, TR=100ms

B_1_^+^ maps were created using both Classic and GLM methods and compared to the reference map. Histograms of the relative difference in B_1_^+^ estimates were calculated for each RF spoiling condition and processing method.

#### ∎ *In vivo* experiments

##### *In vivo* Experiment

Three healthy participants (2 males, 28-40 years) were scanned on the Prisma scanner. Five datasets were acquired per participant, with TR=35ms unless otherwise stated:

1. Interleaved, with *ϕ_BaseInc_* = 120° and TR=100ms. This produced the reference B_1_^+^ map.
2. Interleaved without RF spoiling
3. Interleaved with *ϕ_BaseInc_* = 120°
4. Interleaved with *ϕ_BaseInc_* = 90°
5. Sequentially with *ϕ_BaseInc_* = 120°

Two B1^+^ maps were estimated for each scan using the Classic and GLM methods respectively. The percent difference in B_1_^+^ was calculated with respect to the reference map for the same processing method. To compare processing methods, the percent difference between the two B_1_^+^ maps derived from the reference acquisition was also computed, with respect to that obtained with the Classic method.

Table 1 lists sequence parameters of all experiments.

**Table 1:**
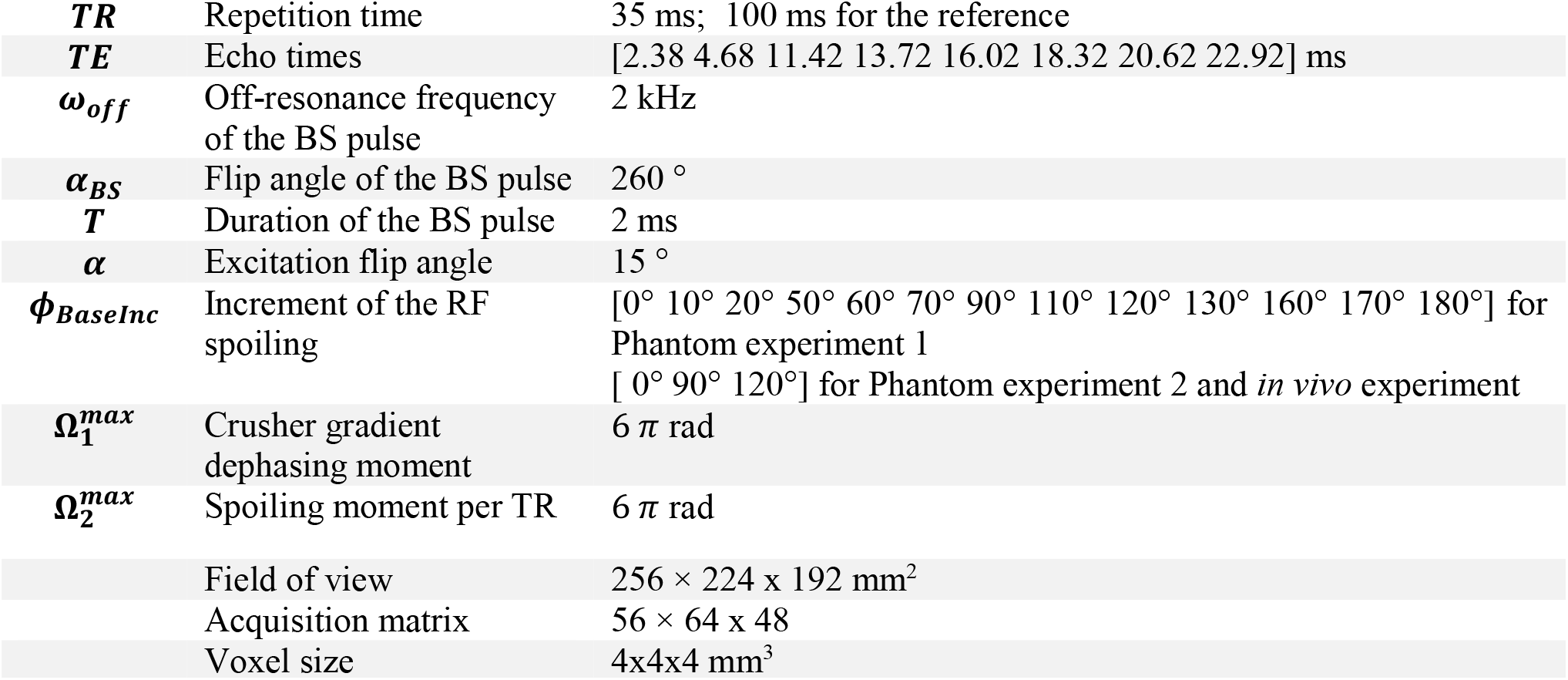
List of the parameters of the phantom and in vivo acquisitions

## Results

### Numerical simulations

#### Effect of acquisition order and RF spoiling

For <u>sequential</u> acquisition ordering, once steady-state was reached for each off-resonance frequency blocks (Fig.2 a), the phase difference between these was zero before the BS pulse (Fig.2 c). It was non-zero after the BS pulse (Fig.2 e) since it contained the BSS phase. However, this BSS phase estimate remained constant over time and was independent of the RF spoiling condition.

**Figure 2:**
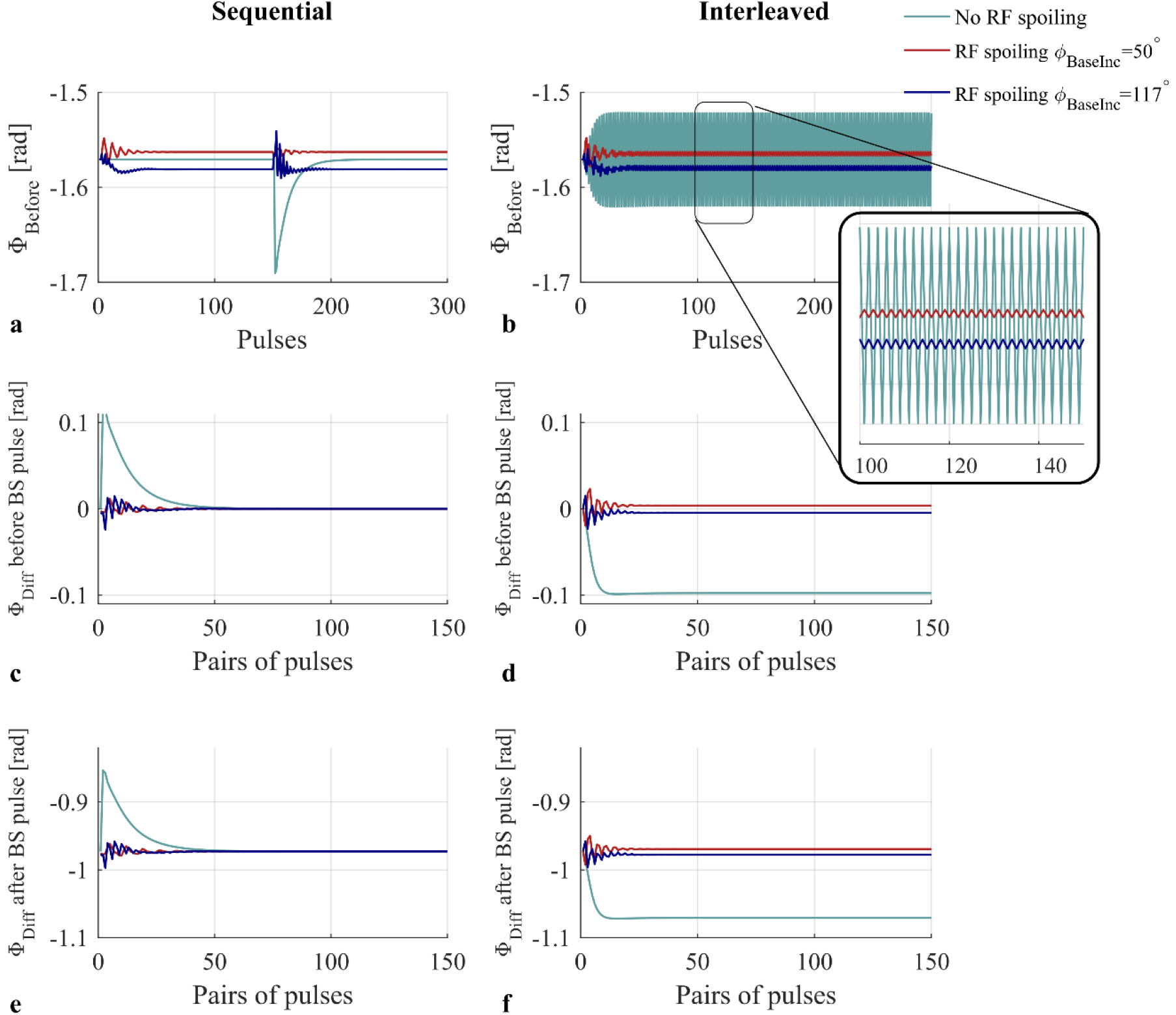
Numerical simulation results. Phase acquired prior to the BS pulse in case of sequential acquisitions (a) or interleaved acquisitions (b), in the presence (red and blue) or not (green) of RF spoiling (Φ_BaseInc_ = 50° and 117°). Phase difference prior to the pulses of the two acquisitions with opposite frequencies (c-d) and phase difference after the BS pulse (e-f).

For <u>interleaved</u> acquisitions, the phase before the BS pulse varied from pulse to pulse regardless of pulse number (Fig.2 b). This temporal variance caused a non-zero phase difference between the interleaved acquisitions with opposite BS pulse frequencies, both before (Fig.2 d) and after (Fig.2 f) the BS pulse. Furthermore, the phase difference after the BS pulse, i.e. the estimate of the BSS phase, differed from the phase difference obtained with the sequential approach and depended strongly on the RF spoiling conditions

#### Effect of RF spoiling increment

The error in the BSS phase resulting from simulating an interleaved acquisition and using the Classic method depended strongly on the RF spoiling increment used (Fig.3). The greatest errors were predicted for phase increments of 0° (equivalent to no RF spoiling) and 180°. Large errors were also predicted for phase increments of 60° and 120° whereas no error was predicted at 90°.

**Figure 3:**
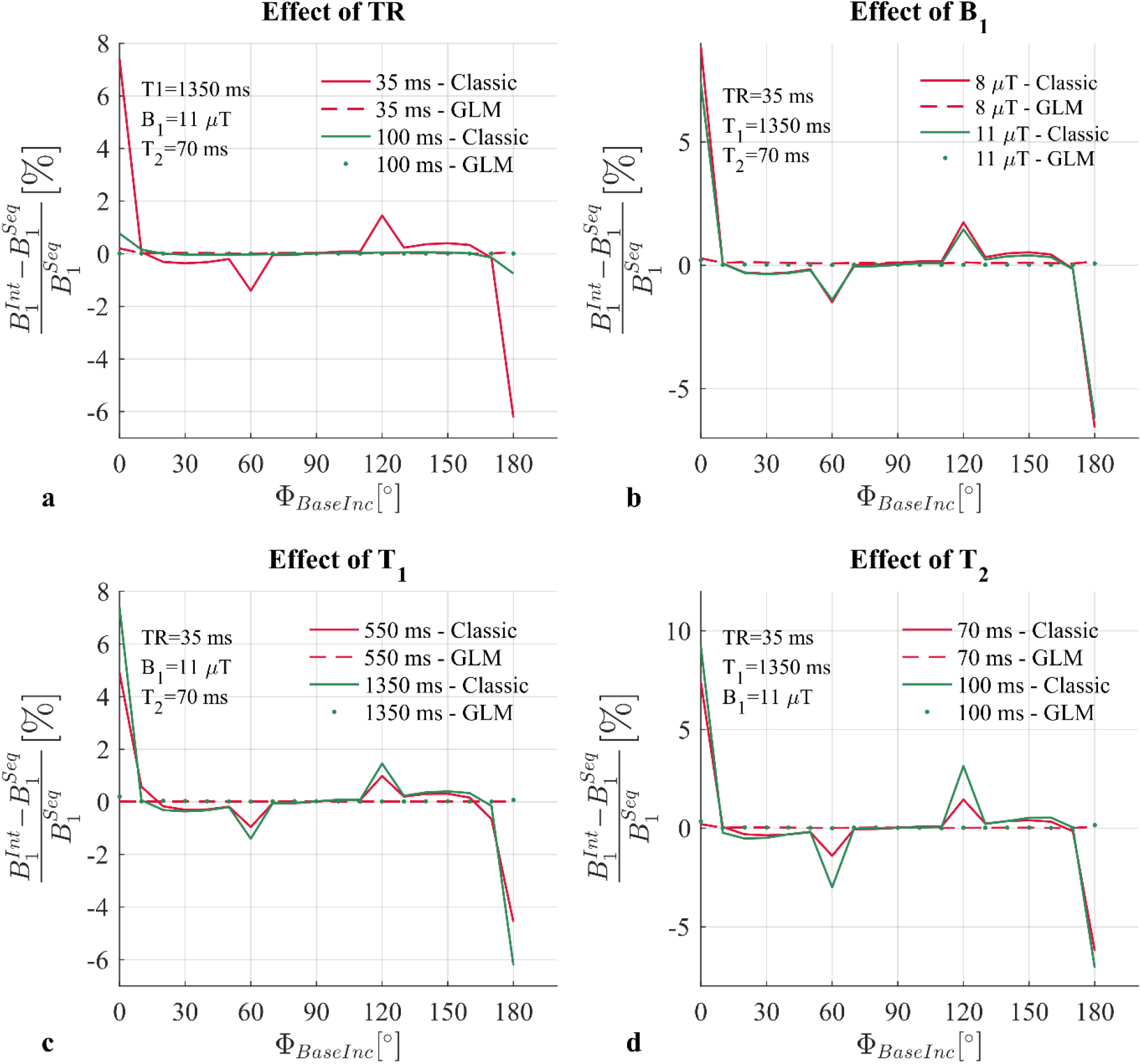
Numerical simulation results. Relative error of the estimate of B_1_^+^ obtained with interleaved acquisitions 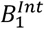 with respect to the estimated B_1_^+^ obtained with sequential acquisitions 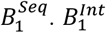 is computed in two ways, the Classic method solid lines, and the GLM method (Eq.4), dashed and dotted lines. 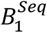 is only computed with the Classic method. The error is observed over a large range of RF spoiling increment (Φ_BaseInc_ ∈ [0:10:180]), for two TR values (a), two B_1_^+^ values (b), two T_1_ values (c) and two T_2_ values (d).

#### Effect of TR, B_1_^+^, T_2_ and T_1_

Increasing TR greatly reduced the predicted error in Φ*_BSS_* (Fig.3 a). Longer T_1_ times led to larger predicted errors, but had less impact than TR, (Fig.3 c). Similarly, longer T_2_ times predicted larger errors (Fig.3 d), especially with phase increments of 60 and 120° where the predicted error was already large.

The amplitude of the BS pulse also had a small impact whereby the *relative* error was predicted to be lower for higher amplitude pulses (Fig.3 b). Note that in this case 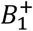 also increases, so while the predicted relative error decreased, the absolute error is actually increased.

#### Impact of GLM method for BSS estimation

The numerical simulations predicted that these errors were removed by using the GLM approach. As a result, the derived B_1_^+^ estimates were predicted to be stable and agree with the B_1_^+^ estimated from the sequential case using the Classic method. This was the case regardless of the RF spoiling conditions, TR, T_1_, T_2_ or BS pulse amplitude.

### Phantom experiments

#### Phantom Experiment 1: Comparing processing approaches

With RF spoiling increments of 0 and 180°, the B_1_^+^ estimates were very different depending on the processing method used (Fig.4), but these estimates agreed when the RF spoiling increment was 90°. Two further peaks in the discrepancy were observed with RF spoiling increments of 60° and 120°. The difference observed without RF spoiling (Φ*_BaseInc_* = 0°) was 3.6%.

**Figure 4:**
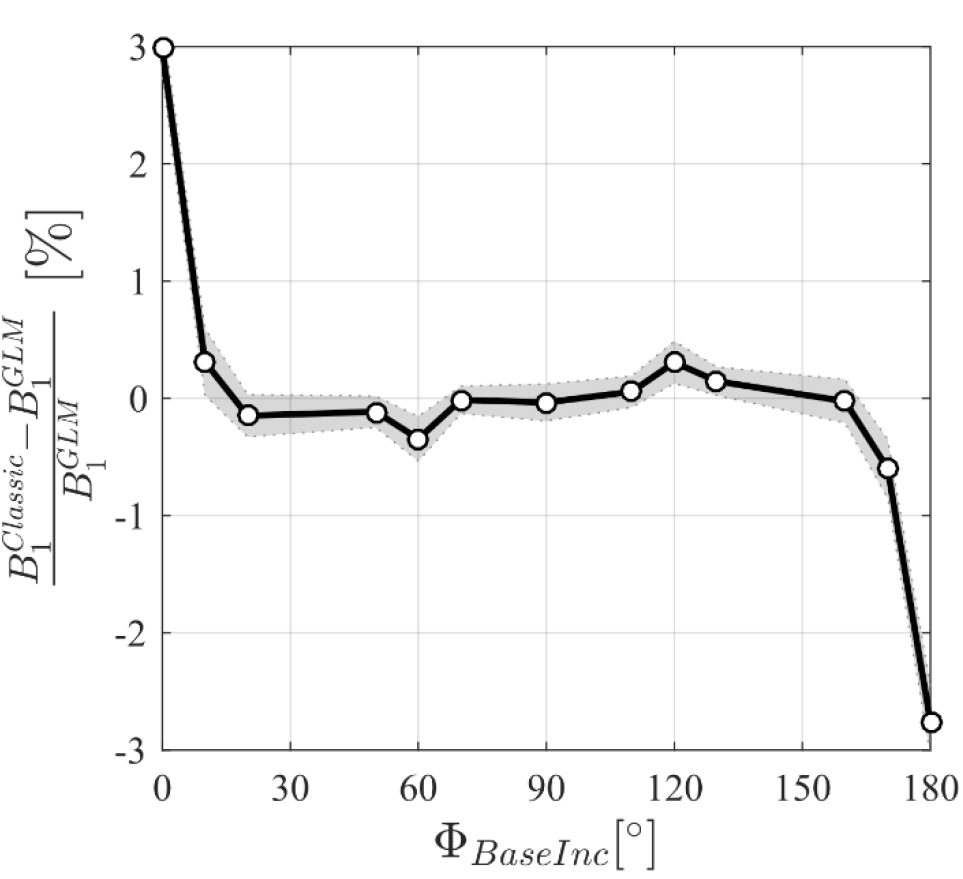
Relative difference of the B_1_^+^ map of a phantom obtained with the Classic method 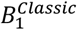, with respect to the B_1_^+^ map reconstructed with the GLM method. One circle is the average difference across the phantom. B_1_^+^ map are obtained with interleaved acquisitions and repeated over a range of RF spoiling increment Φ_BaseInc_ = [0 10 20 50 60 70 90 110 120 130 160 170 180]°.

#### Phantom Experiment 2: Comparing sequential and interleaved approaches

B_1_^+^ maps estimated using the Classic method from interleaved data and sequential data (the reference case) did not agree. The largest bias was observed with no RF spoiling (median (interquartile range, IQR) of 3.04% (0.45%), Fig.5 a). The bias was greatly reduced by RF spoiling with *ϕ_BaseInc_ = 120°* (−0.47% (0.44%)). Negligible bias was observed with *ϕ_BaseInc_ = 90°* (−0.11% (0.36%) or with a longer TR of 100ms and *ϕ_BaseInc_ = 120°* (0.06% (0.34%)).

**Figure 5:**
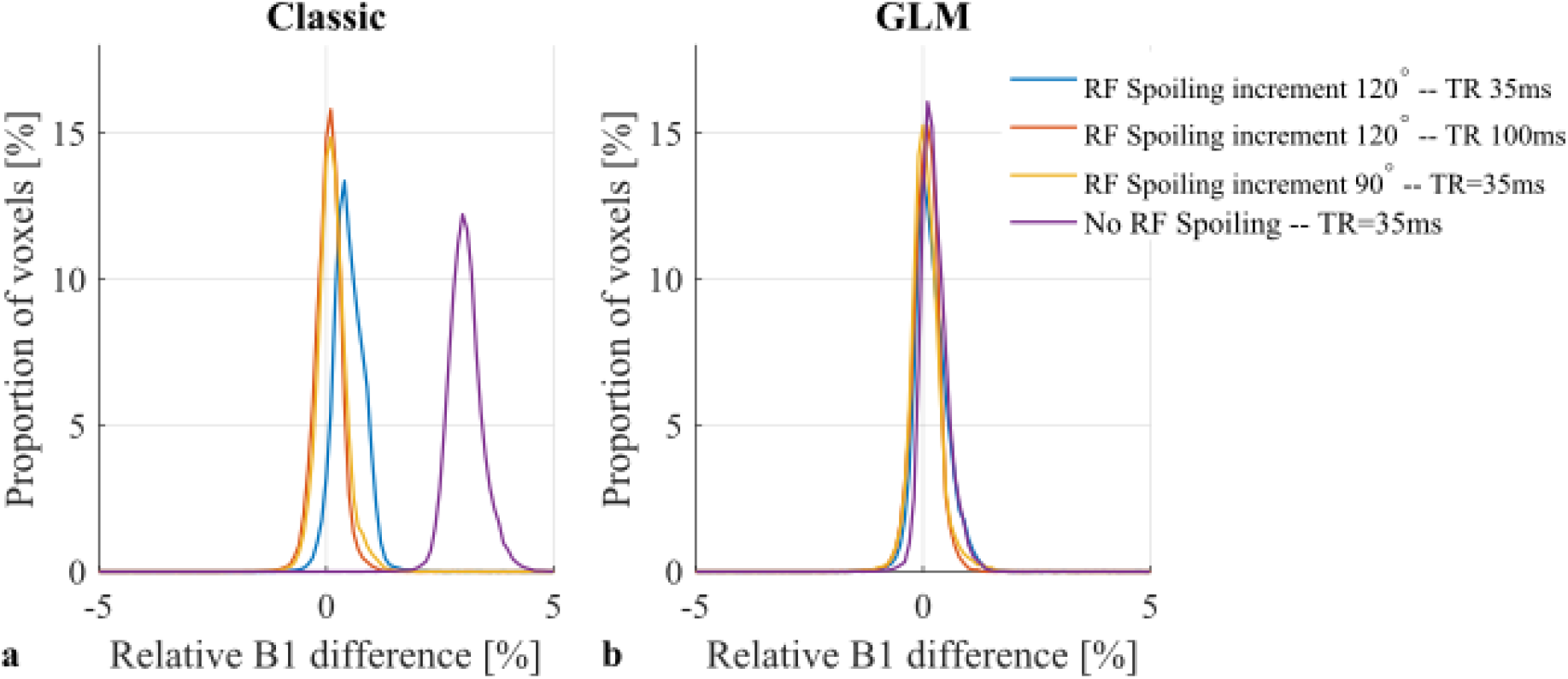
Histograms of the difference in B_1_^+^ measured in a phantom with an interleaved acquisition scheme relative to the reference B_1_^+^ map acquired with a sequential acquisition and processed using the Classic method. The B_1_^+^ maps were calculated using either the Classic (a) or GLM (b) method. The data were acquired with a short TR (35ms) without (purple) or with RF spoiling (Φ_BaseInc_ = 120° (blue) and Φ_BaseInc_ = 90° (yellow)), or a long TR (100ms) and RF spoiling (Φ_BaseInc_ = 90°) (red). The reference B_1_^+^ map, with respect to which the error was calculated, was acquired with sequential ordering and RF spoiling (Φ_BaseInc_ = 90°) and processed using the Classic method.

**Figure 6:**
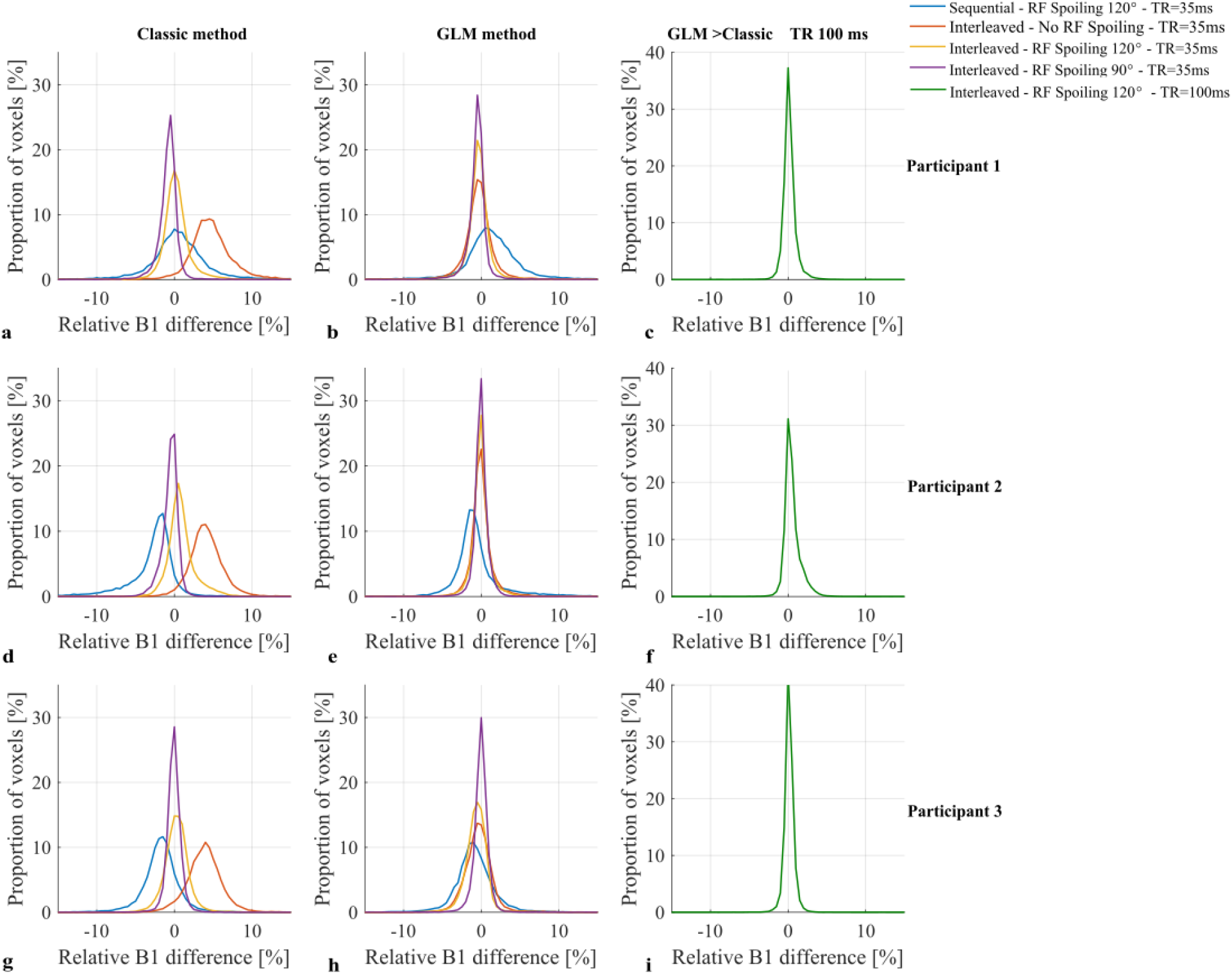
Histograms of the difference in B_1_^+^ relative to the reference B_1_^+^ map acquired in vivo with RF spoiling (Φ_BaseInc_ = 120°) with a TR of 100ms, in interleaved order. The B_1_ maps are either calculated with the classic method (a-d-g) or the GLM method (b-e-h). Percentage difference between the reference B_1_^+^ maps computed with the GLM and the Classic methods (c-f-i). Each row corresponds to a different participant.

The biases observed without RF spoiling and with *ϕ_BaseInc_ = 120°* relative to the sequential case were removed when the GLM method was used to process the same interleaved data (Fig.5 b; 0.21% (0.36%) for no RF spoiling and 0.13% (0.44%) for RF spoiling with *ϕ_BaseInc_* = 120°).

### In vivo experiments

The reference acquisition, which used interleaved ordering, a longer TR of 100ms and an RF spoiling increment of *ϕ_BaseInc_* = 120°, produced consistent B_1_^+^ maps with the Classic and GLM methods (Fig.7 c, f, i). The median (IQR) differences relative to the GLM method were: 0.09% (0.72%), 0.30% (0.97%) and 0.11% (0.60%) for participants 1, 2 and 3, respectively. However, higher B_1_^+^ values were observed in the ventricles of Participant 2 (Fig.8) of the map computed with the classic method. Consistent with this observation, a tail in the histogram was also present for Participant 2 (Fig.7 f).

**Figure 7:**
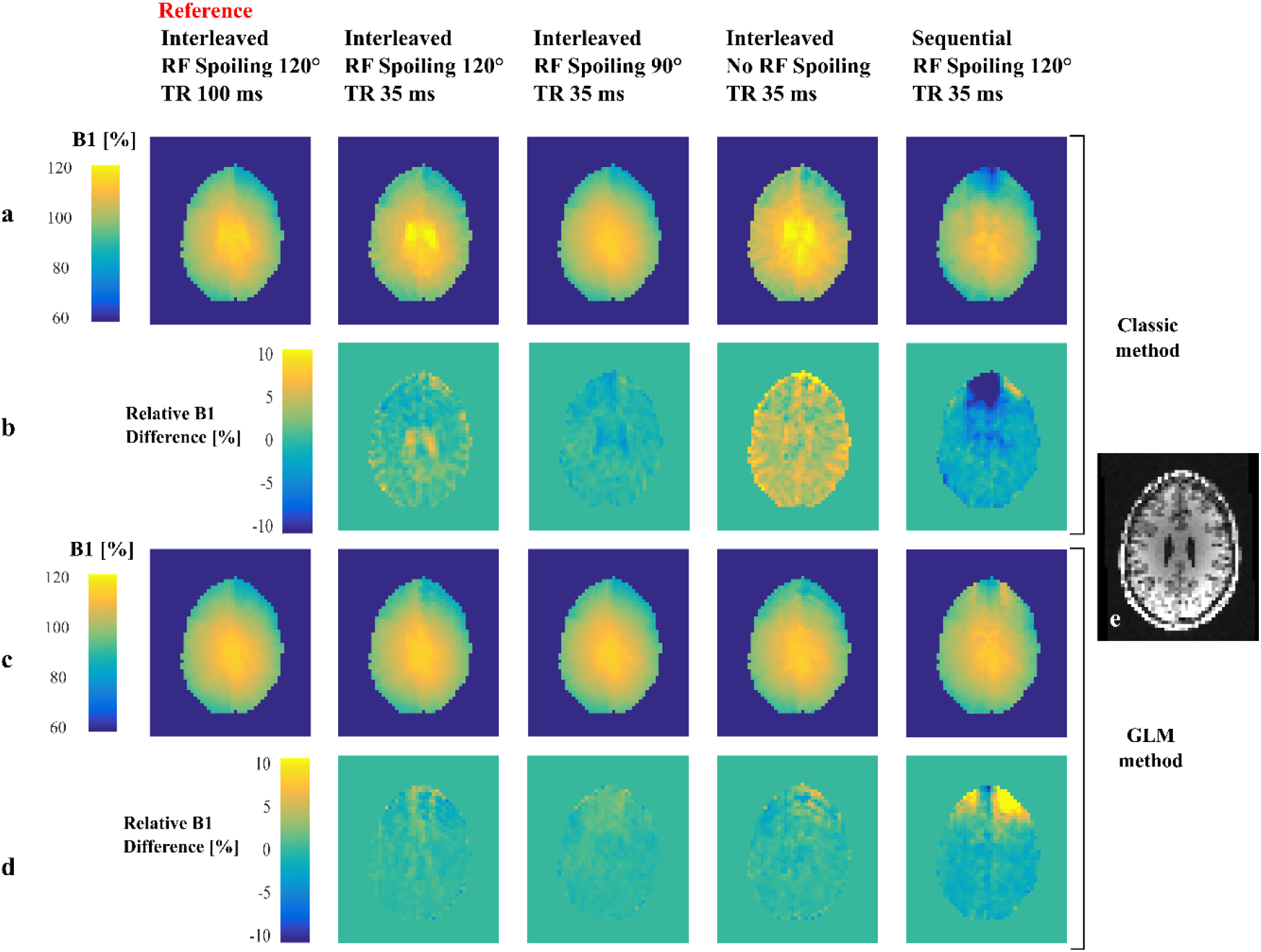
B_1_^+^ maps of one slice obtained on Participant 2 with the Classic (a) and the GLM approach (c) with 5 different protocols. Maps of the relative difference of each B_1_^+^ map with the reference maps computed with the same method: Classic (b) or GLM (d) approach. **e**. Structural image of the same slice, acquired independently.

**Figure 8:**
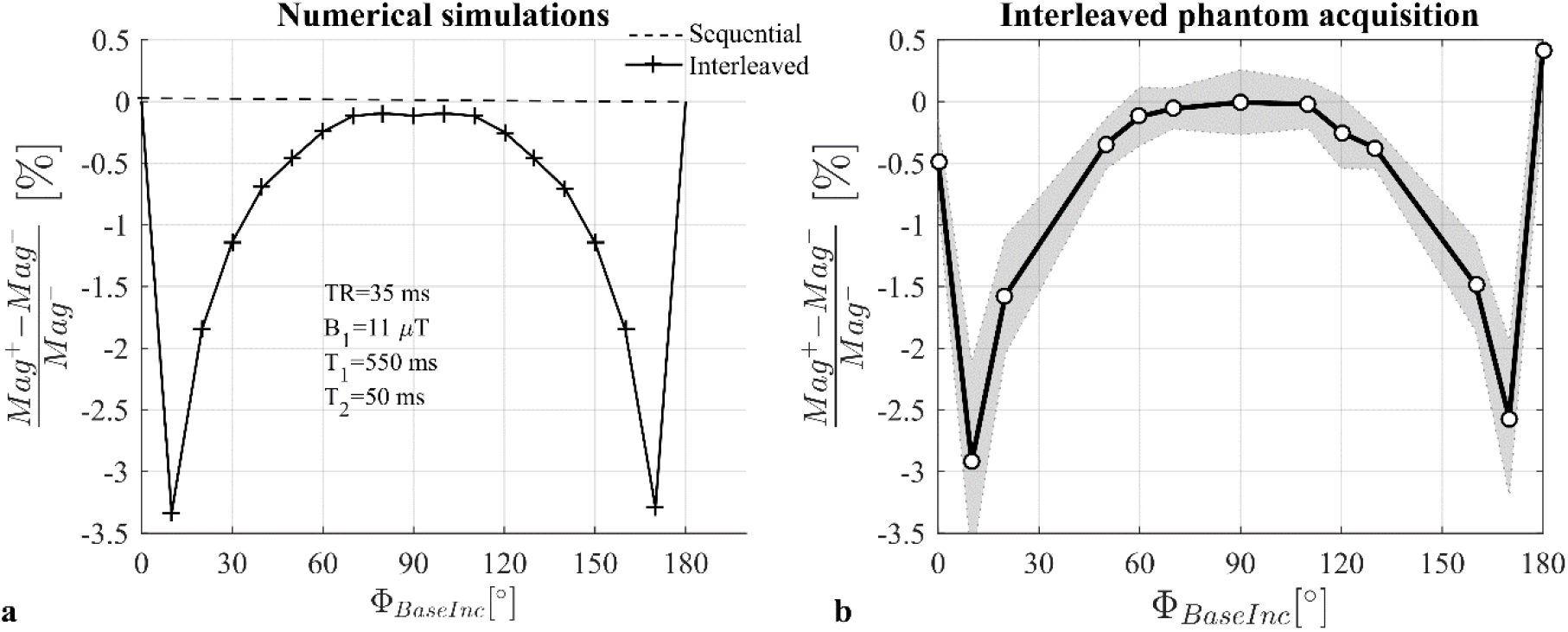
Relative difference in magnitude between the two acquisitions with opposite off-resonance frequencies in case of Interleaved or Sequential order predicted by numerical simulations (c). Relative difference in magnitude between the two interleaved measured on data of Phantom Experiment 1.

Large bias was seen with respect to this reference when B_1_^+^ maps were estimated from interleaved data, acquired without RF spoiling, using the Classic method (Fig.7 a, d, g). Qualitatively, the bias was highly visible in the B_1_^+^ maps (Fig.8 a) following anatomical detail, and was greatest in the ventricles with long T_1_ and T_2_. In the difference map, strong bias was visible along the cortical ribbon (Fig.8 b). Median (IQR) differences were 4.52% (2.92%), 3.99% (2.48%), 3.86% (2.63%) for participants 1, 2 and 3, respectively. These biases were greatly reduced when the data were processed using the GLM method: −0.33% (1.78), −0.1% (1.22) and −0.35% (1.98), respectively (Fig.7 b, e, h; Fig.8 c, d).

The bias was greatly reduced when RF spoiling was used (Φ*_BaseInc_* ∈ [90°, 120°]), and never exceeded 0.69%. However, systematically higher B_1_^+^ values were observed with Φ*_BaseInc_* = 120° compared to Φ*_BaseInc_* = 90° (Fig.7, yellow and purple curves). This difference in B_1_^+^ values was greatly reduced when using the GLM method.

High variability in B_1_^+^ bias was observed when the sequential acquisition ordering was used (Fig.7, blue curves) and artefacts were visible in the B_1_^+^ and difference maps (Fig.8 a-d). This was the case regardless of the processing approach.

## Discussion

Efficient methods for mapping the B_1_^+^ transmit field with high accuracy and precision are prerequisite for demanding MRI applications, such as the quantification of the longitudinal relaxation rate^3^. Biases in B_1_^+^ estimates may underlie inter-site differences in relaxation rates^24^ while uncertainty in the estimates will lower reproducibility^4^. B_1_^+^ mapping based on the phase accrued due to the Bloch-Siegert shift has been reported to be an efficient technique for accurately estimating the spatial distribution of B_1_^+^ when compared to other magnitude or phase-based techniques^7,25^.

Our numerical simulations indicate that the sequential approach for acquiring the necessary BSS data will deliver a bias-free estimate of the B_1_^+^ field. However, in agreement with previous reports^17,10^, our *in vivo* experiments show that this approach is sensitive to phase perturbations over time such as those caused by motion and scanner drifts. The resulting B_1_^+^ maps had visible artefacts and large biases (Fig.8, right column). The greater robustness of the interleaved acquisition scheme has led to its adoption in more recent work using this technique^11,26^.

In the interleaved case, however, our numerical simulations showed that the phase never reached steady-state but rather a pseudo-steady-state that alternated between two conditions depending on the frequency of the preceding off-resonance BS pulse. As a result, the difference in phase between interleaves is not solely due to the BSS effect and therefore does not match the phase difference of the sequential ordering scheme (Fig.2). The additional phase accrued biased the estimated BSS phase (Fig.3) and therefore the B_1_^+^ estimates in phantom and *in vivo* experiments. We have shown that the bias depends on intrinsic tissue properties (T_1_, T_2_) as well as sequence parameters (TR, RF spoiling increment, amplitude of the BS pulse).

Although the biases observed here are relatively small, their impact on the R_1_ estimate can be far greater. For example, the Variable Flip Angle (VFA) technique, widely used for whole brain R_1_ mapping^27–30^ is highly sensitive to B_1_^+^ inhomogeneity and therefore requires correction. In this case, the accuracy of the B_1_^+^ estimate is crucial since it can be shown that a given bias in B_1_^+^ will lead to a bias in the R_1_ estimate that is at least twice as large and increases by even more for acquisitions with high flip angles or large error in the B_1_^+^ estimate. In fact under certain conditions, such as in CSF where the T_1_ and T_2_ are long, the bias can reach 8% (Fig.8) when no RF spoiling is used, which would lead to a minimum of 16% bias in R_1_ with the VFA technique. Hence even small errors must be accounted for if accurate and robust R_1_ estimates are to be obtained.

Here, we have proposed and validated a novel acquisition and processing scheme for interleaved BSS-based B_1_^+^ mapping that does not suffer from these biases. Crucially, multiple echoes are acquired either side of the BS pulse and a GLM framework is used to describe the phase evolution over time. The GLM models the effects of the BS pulse, B_0_ inhomogeneity, eddy currents and phase offsets both common to, and specific to, the positive and negative off-resonance frequency interleaves. The bias observed with the Classic method results from the invalid assumption that the only difference between the two interleaves is the phase imparted by the BS pulse. Indeed, as demonstrated by the numerical simulations, a difference is already present before playing out the BS pulse (Fig.2 d). The use of two distinct regressors (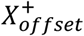 and 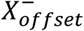) in the model matrix of the GLM allows the two interleaves to differ, even prior to the BS pulse. This removes the bias in B_1_^+^ that would otherwise be present. Numerical simulations (Fig.3) and phantom experiments (Fig.4) confirm this, with both showing peak differences for RF spoiling increments of 0° (equivalent to no RF spoiling), 60°, 120° and 180°. Inversion recovery and multi-echo spin echo experiments indicate T_1_ and T_2_ times of 550 ms and 70 ms for the FBIRN phantom^23^ used. However, the latter estimation did not incorporate any correction for stimulated echoes^31^, and the T_2_ may be as short as 50 ms. Simulations using the same sequence parameters as the phantom experiments, with a T_1_ of 550 ms, and a T_2_ of 70 ms, predicted an error of 4.9% (Fig. 3d) for the case of no RF spoiling. With a shorter T_2_ of 50ms, a lower error of 3.4% was predicted by the simulations (data not shown). These results are in broad agreement with the somewhat lower error of 3.0% observed experimentally for this case. Of note, incorporation of diffusion effects into the simulations^32,33^ had little impact on the level of bias in the estimated B_1_^+^. Furthermore, while it is the phase component that is key to estimating B_1_^+^, it is also worth noting that our simulations predicted a difference in the magnitude of the magnetisation between TRs with interleaved off-resonance frequencies, and that this difference would depend on the RF spoiling increment. Good agreement was again seen between prediction (Fig.8a) and experiment (Fig.8b). No such difference was predicted for sequential ordering of the off-resonance frequencies. Also in agreement with the numerical simulations, the proposed GLM method removed the dependence of the B_1_^+^ estimates on the RF spoiling increment, the TR and the acquisition mode in both phantom (Fig.5) and *in vivo* (Fig.6 and Fig.7) experiments.

The robustness of the GLM method to the sequence parameters, and the RF spoiling increment in particular, makes this method more flexible, which can be exploited to optimize the Signal-to-Noise Ratio (SNR). Given the dependence of the signal amplitude on the RF spoiling increment^34^, a small gain in reproducibility can be expected by choosing the optimal value. In fact, theoretical analysis of the variance of the B_1_^+^ estimates can be used to show that the GLM should deliver higher precision. This has been verified empirically (data not shown) when using the same data for each processing method, as has been done for all of the experiments presented in this work. Although this does not affect the accuracy, it does penalise the Classic approach from a precision perspective since the TE is longer than necessary. Nonetheless, theoretical analysis would also predict improved precision via the GLM when compared to the Classic approach even with an optimal, shorter, TE. Determining the sequence settings that maximise the reproducibility and quantifying the full benefit that can be gained empirically will be the focus of future work.

In theory, the GLM method could use just a single off-resonance frequency with a reduced model matrix containing only half-length regressors for *X_BSS_*, *X*_*ω*_*B*_0___, *X_Even/Odd_* and 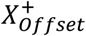^35^. In this case, *X_BSS_* would model all the phase imparted by the BS pulse, including the component depending on B_0_ inhomogeneity and chemical shift. However this could be corrected with the information captured by the second regressor *X*_*ω*_*B*_0___. However, since the BSS phase is imparted only once, the BS pulse flip angle would need to be doubled to achieve the same phase-to-noise ratio. This is problematic from a SAR perspective, and the benefit of a single off-resonance frequency acquisition would be negated if the TR were also doubled to address it. Besides, removing the second acquisition with opposite off-resonance frequency prevents the isolation of the BS phase of interest because any phase caused by the eddy currents of the crushers, for example, would also be captured by the same regressor making the problem ill-posed. A workaround consisting of adding further gradients after the 4^th^ echo, to distinctly capture the effects of eddy currents, has been proposed and tested but has proven to be effective only in phantom experiments^32^.

### Limitations

Given that the GLM method relies on a multi-echo sequence, additional pre-processing steps are required compared to the Classic approach. Phase unwrapping across echoes is necessary. It has been necessary to spatially unwrap the phase difference between successive echoes to deal with large phase accumulation between successive echoes, then cumulatively add these to the first echo.

In the proposed method, multiple echoes are used to estimate the BSS phase, some of the echoes may suffer from dropout and potentially introduce noise into the estimate. To minimize this effect a weighted least-square (WLS) approach had been used to estimate the parameters of the GLM, down-weighting echoes with lower magnitude.

Conventionally, the BS pulse is applied just after the excitation pulse. Here, two echoes preceded the BS pulse and the difference of the third echoes, from the different off-resonance frequency acquisitions, was used to estimate B_1_^+^ using the Classic method. This increases the minimum TE (by ~4ms) and therefore lowers the SNR relative to the single-echo method. However, while this might reduce precision, it would not be expected to introduce bias.

This study focused on short TR 3D acquisitions. For 2D acquisitions, the shot-to-shot inconsistencies may be less problematic since the TR will be longer, although a bias was still observed in long T_1_ regions with a TR of 100 ms (Fig.8– First column). Besides,, the two acquisitions of one slice will be more separated in time which may result in additional phase differences due to motion, similar to the problem affecting sequential acquisitions.

Although more efficient pulses have been proposed^18,36^, only the commonly used Fermi shape for the BS pulse was investigated here. However it can be shown that for the same imparted Bloch-Siegert phase and the same off-resonance frequency, the bias introduced by the interleaved acquisition order is equivalent for a Fermi pulse and the more optimised pulse design suggested by Duan *et al*^18^.

While we have shown how to more accurately estimate the BSS phase, the conversion to 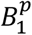 may still be a source of inaccuracy if any assumptions underlying Eq.4 are violated^18^. For the particular conditions we have explored (a Fermi pulse with 2ms duration, *γB*_1_/*ω_off_* = 0.23 and *B*_1_ = 11*μT* the error from this approximation is estimated from simulation to be less than 1%. Regardless of how the phase is converted to a B_1_^+^ value, it is imperative that the bias caused by interleaving the off-resonance pulses be removed.

The precision and accuracy of the GLM technique and the Classic method with ϕ*_BaseInc_* = 90° relative to other B_1_^+^ mapping methods remain to be investigated. However, determining absolute accuracy will always be challenging since every method will have its own limitations.

## Conclusion

Interleaved acquisitions are recommended for Bloch-Siegert based B_1_^+^ mapping to increase robustness to motion and scanner drift. However we have shown that with the Classic estimation method, this can introduce error into the B_1_^+^ estimates that will depend on tissue properties and sequence settings. In theory, one could use an RF spoiling increment of 90° to be immune to this error. However, we have also proposed and validated a multi-echo sequence design, combined with a GLM framework, to robustly isolate the BSS-induced phase regardless of the sequence parameters used. This allows bias free, low error estimates of the B_1_^+^ efficiency that do not depend on tissue properties, sequence settings and would furthermore be immune to reproducible hardware imperfections. Importantly, the proposed modifications do not extend acquisition time, reduce sensitivity or increase SAR. The latter is particularly important since SAR is a limiting factor at higher field strengths.

## Acknowledgment

The Wellcome Centre for Human Neuroimaging is supported by core funding from Wellcome [203147/Z/16/Z].

## Supporting Information

### Bloch-Siegert phase shift expression

The first-order Taylor approximation of the analytical solution of the Bloch-Siegert shift is derived in this section in the same vein as (Duan et al, 2013)^1^

In a static magnetic field, *B*_0_, the precessional frequency is the Larmor frequency given by *ω*_0_ = *γB*_0_. The BSS approach applies an off-resonance RF pulse with a rotating frequency of *ω*_2_, such that *ω_off_* = *ω*_0_ − *ω*_2_ ≠ 0. The off-resonance pulse (termed the BS pulse) modifies the precessional frequency during the time of its application. It has an amplitude *B*_1_ such that *ω*_1_ = *γB*_1_.

In the frame of reference rotating at *ω*_0_, the precessional frequency during this pulse changes to:

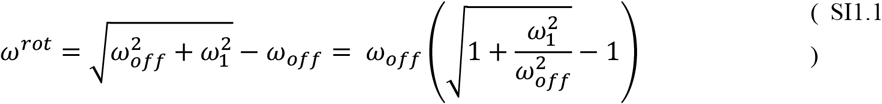

If the off-resonance frequency of the BS pulse is much larger than its strength in frequency units, i.e. *ω_off_* ≫ *ω*_1_ the expression reduces to:

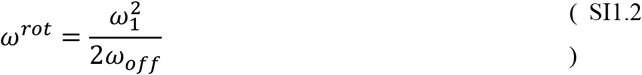

Given that the precessional frequency is no longer equal to *ω*_0_ during the BS pulse, an additional phase is accumulated. Since it is proportional to 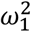 this phase can be used to map the spatial distribution of B1^+^.

Allowing for local *B*_0_ inhomogeneity, the local precessional frequency will be *ω*_0_ + Δ*ω*_*B*_0__. In the rotating frame of reference and under the similar condition as before, (*ω_off_* + Δ*ω*_*B*_0__) ≫ *ω*_1_, the precessional frequency *ω^rot^* during the BS pulse becomes:

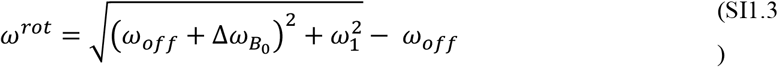

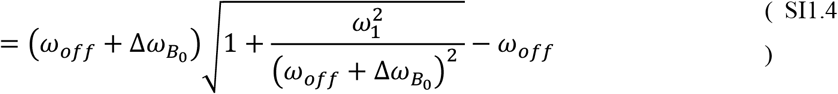

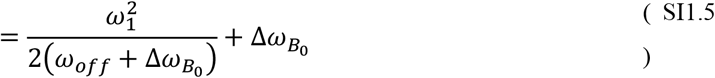

The first term on the right hand side of this expression, representing the frequency shift due to the BS pulse, will be called *ω_BSS_*. Note that this effect adds to the phase accrued due to local *B*_0_ inhomogeneity. The phase specifically accumulated due to the BS pulse, of arbitrary shape and duration T, is given by:

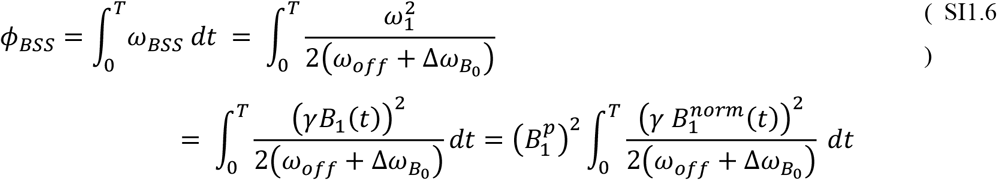

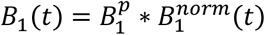, where 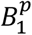 is the peak amplitude of the BS pulse, and 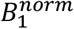 is its normalized shape. Assuming |Δ*ω*_*B*_0__ | ≪ |*ω_off_*|, the first order Taylor expansion of this expression is:

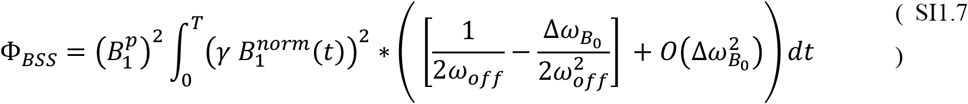

### Numerical simulations

Numerical simulations were carried out to evaluate the phase difference between two acquisitions with opposite off-resonance frequencies both before and after the BS pulse. A typical gradient echo acquisition was simulated in MATLAB (The MathWorks, Inc., Natick, MA) by a series of matrix operations as described below:

1- The magnetization *M* = [*m_x_ m_y_ m_z_*] of a spin ensemble is subject to a rotation by the excitation flip angle *α* about an axis defined by the phase *ϕ* of the excitation pulse.

2- During a subsequent delay TE, longitudinal and transverse relaxation were simulated with time constants *T*_1_ and *T*_2_, respectively. The phase at time TE is termed 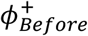 if the off-resonance frequency of the upcoming BS pulse is positive, or 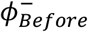 if the off-resonance frequency of the upcoming BS pulse is negative.

3- The application of a crusher gradient rotates the magnetization vector around the z-axis by an angle Ω_1_.

4- The BS pulse, in this work a Fermi pulse described by 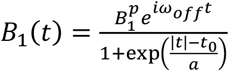, was then simulated as a series of small rotations indexed by *k* ∈ [1:*K*]. Each small rotation k is a rotation of an angle |*B*_1_(*k* * *dt*)| * *dt* about an axis defined by an azimuthal angle of *k* * *dt* * *ω_off_* + *ϕ*.

5- A rotation of the resulting magnetization vector about the z-axis by an angle – Ω_1_ is then applied to simulate the crusher of opposite polarity. The resulting phase at this time is called 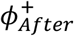 if the frequency of the preceding BS pulse was positive, or 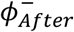 if the frequency was negative. For simplicity, the pulse and crushers were assumed to be instantaneous from a relaxation standpoint such that Φ_*After*_ and Φ_*Before*_ have the same TE. Note that the relaxation is instead accounted for in step 7.

6- Following the second crusher gradient, a spoiler gradient was applied such that the magnetization vector from step 5 was rotated around the z-axis by an angle of *Ω*_2_.

7- T_1_ and T_2_ relaxation were then simulated for the remainder of the TR period, i.e. TR-TE.

8- When RF spoiling was simulated, the phase of the RF pulses, *ϕ*, was incremented such that *ϕ* = *ϕ* + *ϕ_inc_* and ϕ_*inc*_ = ϕ_*inc*_ + ϕ_*BaseInc*_.

These steps were repeated *N_exc_* times, by updating the magnetization of Step 1 with the magnetization vector resulting from Step 7.

The simulation was performed for a 2D grid of spin ensembles (*N_spin_* * *N_spin_*) with varying values for Ω_1_ and Ω_2_ ranging from 0° to 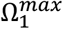 and 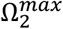, respectively. The values of the crusher dephasing Ω_1_ varied along one dimension of the grid, whereas the values of the spoiler dephasing Ω_2_ varied in the other direction to simulate crushers and spoiler along orthogonal axes. The integral of these spins gave the magnetisation in a single voxel.

For interleaved ordering, the sign of *ω_off_* was switched before each BS pulse. For sequential ordering the sign was only switched after half the total number of pulses 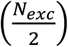. If RF spoiling was included in the simulation, the RF spoiling phase (ϕ and *ϕ_Inc_*) were reset to 0 when the sign of *ω_off_* was switched in the sequential case. In the case of an interleaved acquisition, the phase increment occurred for each repetition, including when the off-resonance frequency also changes.

The Matlab code is available here: https://github.com/fil-physics/Publication-Code/tree/master/Bloch-Siegert.

The numerical values of the parameters used in the simulations are listed in the Supporting Information Table S1

**Supporting Information Table S1:** Parameters used in the numerical simulations

|  |  |  |
| --- | --- | --- |
| $T_1$ | Longitudinal relaxation time | [550, 1350] ms |
| $T_2$ | Transverse relaxation time | [70,100] ms |
| $TR$ | Repetition time | [35, 100] ms |
| $TE$ | Echo time for $\Phi_{Before}$ and $\phi_{After}$ | 2 ms |
| $\alpha$ | Excitation flip angle | 15° |
| $\omega_{off}$ | BS pulse off-resonance frequency | 2 kHz |
| $B_1$ | BS pulse amplitude | [8, 11] $\mu T$ |
| $T$ | BS pulse duration | 2 ms |
| $\phi_{BaseInc}$ | RF spoiling increment | [0 50 117] ° (Figure 2)<br>[0:10:180] ° (Figure 3) |
| $\Omega_1^{max}$ | Crusher gradient dephasing moment | 8 $\pi$ |
| $\Omega_2^{max}$ | Spoiler gradient dephasing moment | 4 $\pi$ |
| $N_{spin}$ | Number of spins in ensemble along each dimension | 100 |
| $N_{exc}$ | Number of excitation pulses simulated | 300 |
| $a$ | Fermi pulse parameter | 0.16 ms |
| $t_0$ | Fermi pulse parameter | 3 ms |

The BS phase estimated by the Classic approach is simulated as:

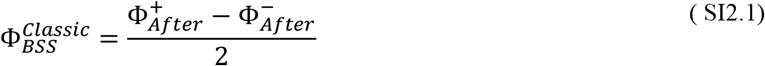

The BS phase estimated by the GLM approach is simulated as:

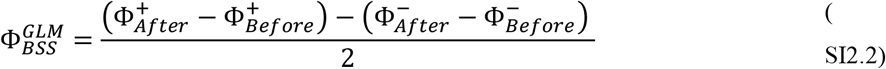

## References

1. Sacolick LI, Sun L, Vogel MW, Dixon WT, Hancu I. Fast radiofrequency flip angle calibration by Bloch–Siegert shift. Magn. Reson. Med. 2011;66:1333–1338 doi: 10.1002/mrm.22902.

2. Padormo F, Beqiri A, Hajnal JV, Malik SJ. Parallel transmission for ultrahigh-field imaging: Parallel Transmission for Ultrahigh-Field Imaging. NMR Biomed. 2016;29:1145–1161 doi: 10.1002/nbm.3313.

3. Lutti A, Dick F, Sereno MI, Weiskopf N. Using high-resolution quantitative mapping of R1 as an index of cortical myelination. NeuroImage 2014;93:176–188 doi: 10.1016/j.neuroimage.2013.06.005.

4. Lee Y, Callaghan MF, Nagy Z. Analysis of the Precision of Variable Flip Angle T1 Mapping with Emphasis on the Noise Propagated from RF Transmit Field Maps. Front. Neurosci. 2017;11 doi: 10.3389/fnins.2017.00106.

5. Ramsey NF. Resonance transitions induced by perturbations at two or more different frequencies. Phys. Rev. 1955;100:1191.

6. Bloch F, Siegert A. Magnetic Resonance for Nonrotating Fields. Phys. Rev. 1940;57:522–527 doi: 10.1103/PhysRev.57.522.

7. Pohmann R, Scheffler K. A theoretical and experimental comparison of different techniques for B1 mapping at very high fields. NMR Biomed. 2013;26:265–275 doi: 10.1002/nbm.2844.

8. Carinci F, Santoro D, von Samson-Himmelstjerna F, Lindel TD, Dieringer MA, Niendorf T. Characterization of Phase-Based Methods Used for Transmission Field Uniformity Mapping: A Magnetic Resonance Study at 3.0 T and 7.0 T. PLoS ONE 2013;8 doi: 10.1371/journal.pone.0057982.

9. Sacolick LI, Wiesinger F, Hancu I, Vogel MW. B1 Mapping by Bloch-Siegert Shift. Magn. Reson. Med. Off. J. Soc. Magn. Reson. Med. Soc. Magn. Reson. Med. 2010;63:1315–1322 doi: 10.1002/mrm.22357.

10. Lesch A, Petrovic A, Stollberger R. Robust implementation of 3D Bloch Siegert B1 mapping. In: Proc. Intl. Soc. Mag. Reson. Med. 23. Toronto; 2015.

11. Lesch A, Schlöegl M, Holler M, Bredies K, Stollberger R. Ultrafast 3D Bloch–Siegert B-mapping using variational modeling. Magn. Reson. Med. 2019;81:881–892 doi: 10.1002/mrm.27434.

12. Saranathan M, Khalighi MM, Glover GH, Pandit P, Rutt BK. Efficient bloch-siegert B1+ mapping using spiral and echo-planar readouts. Magn. Reson. Med. 2013;70:1669–1673 doi: 10.1002/mrm.24599.

13. Basse-Lüsebrink TC, Kampf T, Fischer A, et al. SAR-reduced spin-echo-based Bloch–Siegert B1+ mapping: BS-SE-BURST. Magn. Reson. Med. 2012;68:529–536 doi: 10.1002/mrm.23259.

14. Basse-Lüsebrink TC, Sturm VJF, Kampf T, Stoll G, Jakob PM. Fast CPMG-based Bloch-Siegert B1+ mapping. Magn. Reson. Med. 2012;67:405–418 doi: 10.1002/mrm.23013.

15. Sacolick LI, Lee SK, Grissom WA, Vogel MW. Fast Spin Echo Bloch-Siegert B1 Mapping. In: Proc. Intl. Soc. Mag. Reson. Med. 19. Montreal; 2011.

16. Sturm VJF, Basse-Lüsebrink TC, Kampf T, Stoll G, Jakob PM. Improved encoding strategy for CPMG-based Bloch-Siegert B mapping. Magn. Reson. Med. 2012;68:507–515 doi: 10.1002/mrm.23232.

17. Kameda H, Kudo K, Matsuda T, et al. Improvement of the repeatability of parallel transmission at 7T using interleaved acquisition in the calibration scan. J. Magn. Reson. Imaging 2018;48:94–101 doi: 10.1002/jmri.25903.

18. Duan Q, van Gelderen P, Duyn J. Improved Bloch-Siegert Based B1 Mapping by Reducing Off-Resonance Shift. NMR Biomed. 2013;26:1070–1078 doi: 10.1002/nbm.2920.

19. Bernstein MA, King KF, Zhou XJ. Handbook of MRI Pulse Sequences.; 2004. doi: 10.1016/B978-0-12-092861-3.X5000-6.

20. Hansen MS, Sørensen TS. Gadgetron: an open source framework for medical image reconstruction. Magn. Reson. Med. 2013;69:1768–1776 doi: 10.1002/mrm.24389.

21. Walsh DO, Gmitro AF, Marcellin MW. Adaptive reconstruction of phased array MR imagery. Magn. Reson. Med. 2000;43:682–690 doi: 10.1002/(SICI)1522-2594(200005)43:5<682::AID-MRM10>3.0.CO;2-G.

22. Abdul-Rahman HS, Gdeisat MA, Burton DR, Lalor MJ, Lilley F, Moore CJ. Fast and robust three-dimensional best path phase unwrapping algorithm. Appl. Opt. 2007;46:6623–6635 doi: 10.1364/AO.46.006623.

23. Glover GH, Mueller BA, Turner JA, et al. Function Biomedical Informatics Research Network Recommendations for Prospective Multi-Center Functional Magnetic Resonance Imaging Studies. J. Magn. Reson. Imaging 2012;36:39–54 doi: 10.1002/jmri.23572.

24. Lee Y, Callaghan MF, Acosta-Cabronero J, Lutti A, Nagy Z. Establishing intra- and inter-vendor reproducibility of T1 relaxation time measurements with 3T MRI. Magn. Reson. Med. 2019;81:454–465 doi: 10.1002/mrm.27421.

25. Park DJ, Bangerter NK, Javed A, Kaggie J, Khalighi MM, Morrell GR. A Statistical Analysis of the Bloch-Siegert B1 Mapping Technique. Phys. Med. Biol. 2013;58 doi: 10.1088/0031-9155/58/16/5673.

26. Weingärtner S, Zimmer F, Metzger GJ, Uğurbil K, Van de Moortele P-F, Akçakaya M. Motion-Robust Cardiac B1+ Mapping at 3T using Interleaved Bloch-Siegert Shifts. Magn. Reson. Med. 2017;78:670–677 doi: 10.1002/mrm.26395.

27. Deoni SCL, Peters TM, Rutt BK. High-resolution T1 and T2 mapping of the brain in a clinically acceptable time with DESPOT1 and DESPOT2. Magn. Reson. Med. 2005;53:237–241 doi: 10.1002/mrm.20314.

28. Lescher S, Jurcoane A, Veit A, Bähr O, Deichmann R, Hattingen E. Quantitative T1 and T2 mapping in recurrent glioblastomas under bevacizumab: earlier detection of tumor progression compared to conventional MRI. Neuroradiology 2015;57:11–20 doi: 10.1007/s00234-014-1445-9.

29. Stikov N, Boudreau M, Levesque IR, Tardif CL, Barral JK, Pike GB. On the accuracy of T1 mapping: Searching for common ground. Magn. Reson. Med. 2015;73:514–522 doi: 10.1002/mrm.25135.

30. Weiskopf N, Suckling J, Williams G, et al. Quantitative multi-parameter mapping of R1, PD*, MT, and R2* at 3T: a multi-center validation. Front. Neurosci. 2013;7 doi: 10.3389/fnins.2013.00095.

31. Lebel RM, Wilman AH. Transverse relaxometry with stimulated echo compensation. Magn. Reson. Med. 2010;64:1005–1014 doi: 10.1002/mrm.22487.

32. Gudbjartsson H, Patz S. Simultaneous Calculation of Flow and Diffusion Sensitivity in Steady-State Free Precession Imaging. Magn. Reson. Med. Off. J. Soc. Magn. Reson. Med. Soc. Magn. Reson. Med. 1995;34:567–579.

33. Weigel M. Extended phase graphs: Dephasing, RF pulses, and echoes – pure and simple. J. Magn. Reson. Imaging 2015;41:266–295 doi: 10.1002/jmri.24619.

34. Preibisch C, Deichmann R. Influence of RF spoiling on the stability and accuracy of T1 mapping based on spoiled FLASH with varying flip angles. Magn. Reson. Med. 2009;61:125–135 doi: 10.1002/mrm.21776.

35. Corbin N, Acosta-Cabronero J, Weiskopf N, Callaghan MF. Rapid B1 mapping based on the Bloch-Siegert shift using a single offset frequency and multi-echo readout. In: Proc. Intl. Soc. Mag. Reson. Med. 26. Paris; 2018.

36. Khalighi MM, Rutt BK, Kerr AB. Adiabatic RF pulse design for Bloch-Siegert B mapping. Magn. Reson. Med. 2013;70:829–835 doi: 10.1002/mrm.24507.

## References

Duan Q, van Gelderen P, Duyn J. Improved Bloch-Siegert Based B1 Mapping by Reducing Off-Resonance Shift. NMR Biomed. 2013;26:1070–1078 doi: 10.1002/nbm.2920.

